# NeuroMTL iSEG challenge methods

**DOI:** 10.1101/278465

**Authors:** Vladimir S. Fonov, Andrew Doyle, Alan C. Evans, D. Louis Collins

## Abstract

We propose a tissue classification method for MRI scans of 6-month old infants, and used it to process the iSeg-2017 challenge data. The method relies on a deep-learning 3D U-Net network, trained with MRI scans of 216 infants, aged 6-24 months, from the ACE-IBIS longitudinal database.

## 1 Introduction

Automatic image processing of infant MRI scans is difficult due to the rapid change of anatomy and MR relaxation parameters of human brain tissues during the first year of life. In particular, due to ongoing myelination, the contrast between white matter (WM) and gray matter (GM) is inverted between birth and 12 months in both T1 and T2 weighted images. This process is gradual and in general follows the maturation pattern from central to peripheral, from inferior to superior and from posterior to anterior [Barkovich 1988].

Previously published papers describe methods where infants’ longitudinal scans were segmented by propagating information about tissue location from later timepoints back to earlier time-points [Feng Shi 2010], but the proposed iSeg-2017 dataset only includes images from one timepoint in ten 6-month old infants. Therefore, it is impossible to directly use this method.

In our approach we combine two techniques: i) We first create an extended training dataset by applying a standard tissue classification technique to scans of 24-month old infants from a separate longitudinal dataset (ACE-IBIS) [Hazlett 2017] and then propagating this segmentation back to the 6-month old scans. ii) We then used deep learning to train a 3D U-Net network to automatically perform tissue segmentation on scans of 6-month old infants first on ACE-IBIS scans and then on iSEG scans.

Resulting cross-validation experiment showed that our method achieves good results in terms of kappa overlap measure.

## 2 Methods and materials

### 2.1 Materials

We used data provided by the challenge organizers (iSEG) and another independent dataset (ACE-IBIS) from [Hazlett 2017]. The iSEG dataset consisted of 10 pairs of pre-processed T1w and T2w scans of 6-month old infants. ACE-IBIS is a longitudinal study comprising 318 infants at high familial risk (HR) for autism spectrum disorder (ASD), and 117 infants at low familial risk (LR) for ASD. Out of this dataset we selected a subset of scans corresponding to subjects that have both T1w and T2w scans acquired at 6, 12 and 24 months (n=216 subjects).

### 2.2 Methods: longitudinal tissue classification of ACE-IBIS scans

We used following process to created segmentation training library based on ACE-IBIS dataset:

i) An unbiased population average of T1w scans for each age group was created using the method described in [Fonov 2010] ii) The group average for 24-month old scans were manually segmented into areas of high probability of GM, WM and CSF. iii) All 24-month old scans were non-linearly registered to the 24-month template using the T1w modality. Tissue priors were then transformed to the space of subject’s scan. iv) An expectation-maximization algorithm was run to obtain tissue classification. v) Longitudinal non-linear registrations between scans at 6 and 12 months and then between 12 and 24 months were performed using ANTs with mutual information [Avants 2011], using both T1w and T2w image modalities. Non-linear transformations from 6 to 12 and from 12 to 24 were concatenated and tissue classification maps from 24 months were transformed to match 6-month old scans.

### 2.3 Methods: deep learning

We adopted a 3D U-Net [5] for tissue classification, with 5 downsampling and 5 upsampling layers. Max-pooling was used, halving the patch resolution in downsampling steps and convolutional kernels that upsample by a factor of 2. Each layer also used batch-normalization and ReLU non-linear activation functions. The final layer of the network contains two 1 × 1 × 1 convolutional layers with 64 and 32 channels and includes a dropout layer. The final layer is a linear mapping of 32 features to the three segmentation labels (GM, WM, CSF) implemented again as 1 × 1 × 1 convolution. The output volume is cropped to remove border voxels to reduce edge effects. The loss function used to train the network was chosen as a log softmax of the categorical cross-entropy, and a generalized kappa overlap metric was used to track performance on an out-of-sample validation dataset. The total number of trainable parameters in the model was 2,810,296. A diagram illustrating the architecture of the network is shown on Fig. 1. The parameters of each layer are shown in Table 1.

**Table 1.** Parameters of U-net

| U-net layer | Input Channels | Output Channels | Convolution kernel 1 | Convolution kernel 2 | Upsampling kernel |
| --- | --- | --- | --- | --- | --- |
| 1 | 4 | 64 | 5x5x5 | 5x5x5 | 5x5x5 |
| 2 | 16 | 64 | 5x5x5 | 5x5x5 | 3x3x3 |
| 3 | 16 | 64 | 3x3x3 | 3x3x3 | 3x3x3 |
| 4 | 16 | 64 | 3x3x3 | 3x3x3 | 3x3x3 |
| 5 | 32 | 64 | 1x1x1 | 3x3x3 | - |

**Fig. 1.**
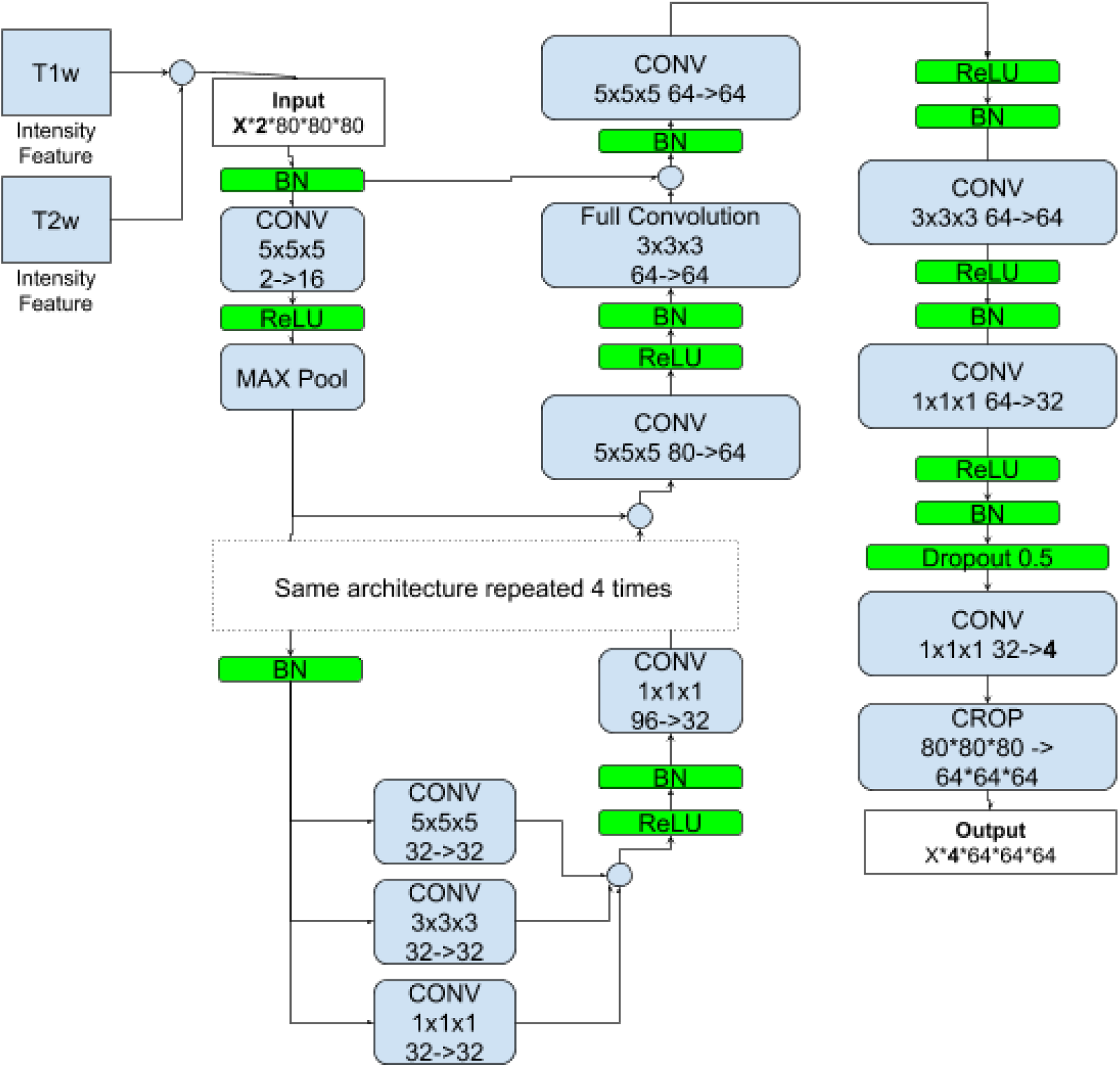
Deep net architecture (U-Net), X correspond to minibatch size (1).

We used both T1w and T2w scans, normalized to have zero mean and unit standard deviation. An Adam optimizer was used for both training stages: the ACE-IBIS training stage used 10000 mini-batches, the iSEG training used 4000 mini-batches, each mini-batch contained a 80 × 80 × 80 voxel patch randomly drawn from ROI around brain. The iSEG data was additionally augmented by including samples that were flipped around Y-Z plane, and also applying small random rotations and displacements. Hyper-parameters were chosen based on performance of an out-of-sample validation dataset. Figure 2 shows the progress of training.

**Fig. 2.**
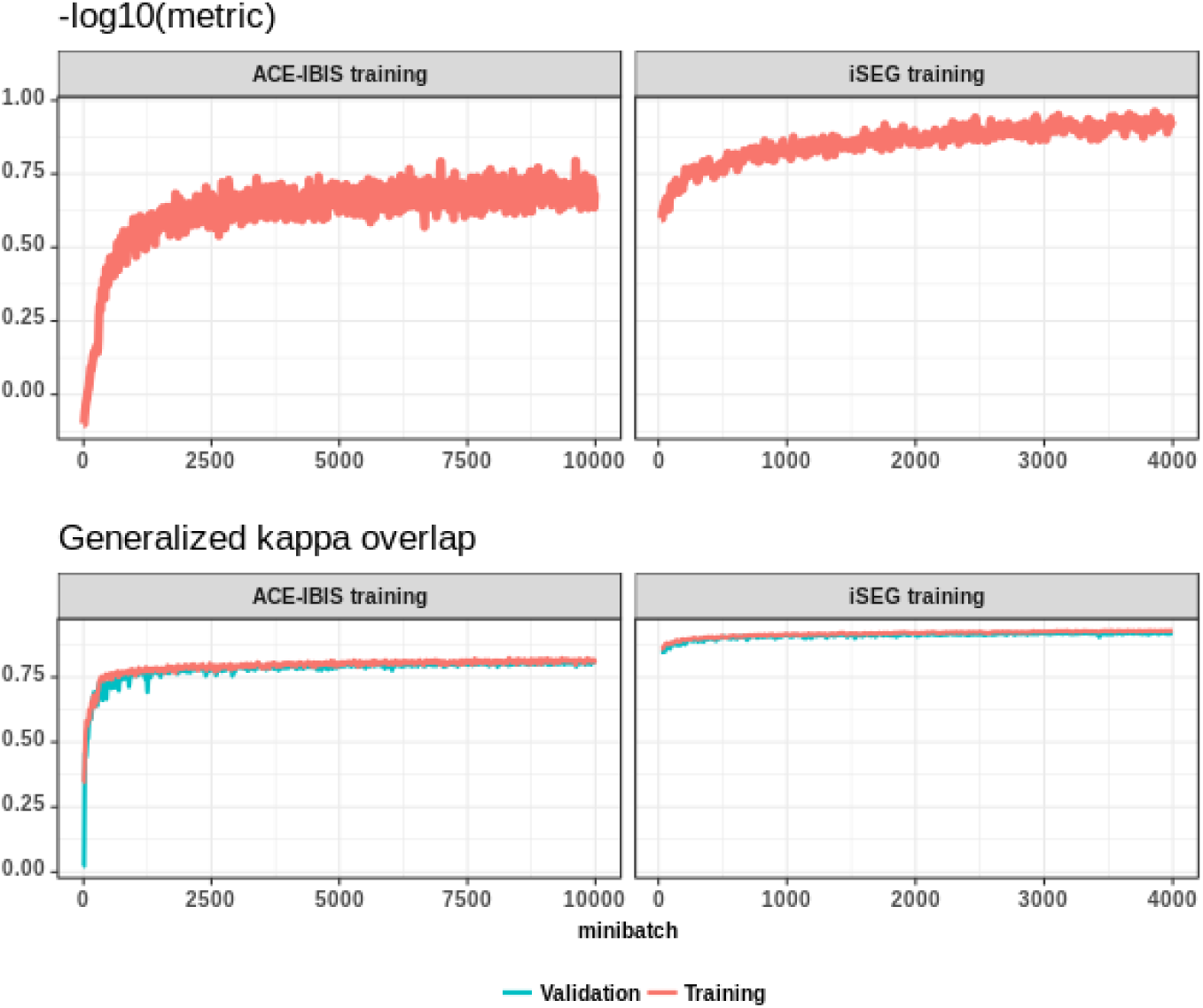
Progress of deep-net training.

## 3 Results

### 3.1 Cross-validation experiment

Hyper-parameters of the deep net were chosen based on leave-one out cross-validation experiment. Results of the final version are shown in Fig. 3. For the application on the testing dataset from iSEG challenge, the deep-net was re-trained using all 10 samples from iSEG training dataset (with data augmentation as described above). The resulting network was then applied to classify testing datasets. All experiments were performed on computer with Xeon CPU E5-2620 v4 @ 2.10GHz with 64GB of ram and Nvidia Titan-X GPU, with deep-net implemented in Torch. Training on ACE-IBIS dataset took approximately 32 hours, final training on iSEG data - 11 hours. Application on a single dataset, using GPU, takes 8 seconds.

**Fig. 3.**
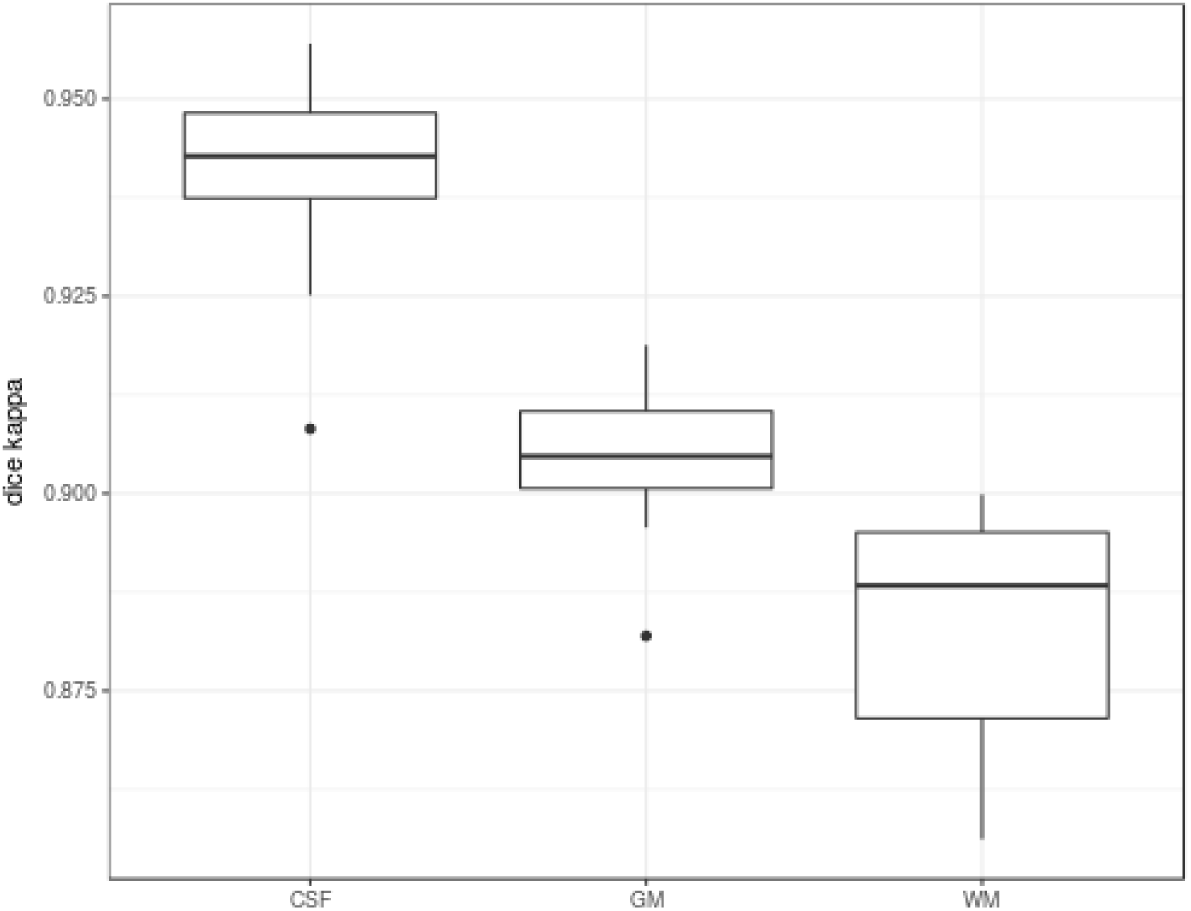
Results of cross-validation.

### 3.2 iSEG challenge results

Deep-net was re-trained on the whole iSEG datasets (without validation subset) and cross-validation, with 4000 minibatches. Trained deep-net was applied to the testing dataset from iSEG web site and sent to the organizers under the name ***NeuroMTL.***

Resulting metrics from http://iseg2017.web.unc.edu/rules/results/ are shown in Table 2, where DICE corresponds to Dice overla metric, MHD - Modified Hausdorff Distance in mm, AHD - Average Surface Distance

**Table 2.** iSEG challenge results (http://iseg2017.web.unc.edu/rules/results/)

|  | CSF |  |  | GM |  |  | WM |  |  |
| --- | --- | --- | --- | --- | --- | --- | --- | --- | --- |
|  | DICE | MHD | ASD | DICE | MHD | ASD | DICE | MHD | ASD |
| Mean | 0.94 | 11.06 | 0.20 | 0.90 | 7.11 | 0.39 | 0.89 | 6.87 | 0.46 |
| Std | 0.02 | 2.67 | 0.06 | 0.01 | 1.92 | 0.05 | 0.02 | 1.25 | 0.07 |

### 3.3 Open source implementation

Scripts used to implement and train deep-net are available at https://github.com/vfonov/NeuroMTL_iSEG

